# The BvgS PAS domain: an independent sensory perception module in the *Bordetella* BvgAS phosphorelay

**DOI:** 10.1101/614891

**Authors:** M. Ashley Sobran, Peggy A. Cotter

## Abstract

To detect and respond to the diverse environments they encounter, bacteria often use two-component regulatory systems (TCSs) to coordinate essential cellular processes required for survival. In pathogenic *Bordetella* species, the BvgAS TCS regulates expression of hundreds of genes, including those encoding all known protein virulence factors, and its kinase activity is essential for respiratory infection. Maintenance of BvgS kinase activity in the lower respiratory tract (LRT) depends on the function of another TCS, PlrSR. While the periplasmic venus fly-trap domains of BvgS have been implicated in responding to so-called modulating signals *in vitro* (nicotinic acid and MgSO_4_), a role for the cytoplasmic Per-Arnt-Sim (PAS) domain in signal perception has not previously been demonstrated. By comparing *B. bronchiseptica* strains with mutations in the PAS domain-encoding region of *bvgS* with wild-type bacteria *in vitro* and *in vivo*, we found that although the PAS domain is not required to sense modulating signals *in vitro*, it is required for the inactivation of BvgS that occurs in the absence of PlrS in the LRT of mice, suggesting that the BvgS PAS domain functions as an independent signal perception domain. Our data also indicate that the BvgS PAS domain is important for controlling absolute levels of BvgS kinase activity and the efficiency of the response to modulating signals *in vitro*. Our results indicate that BvgS is capable of integrating sensory inputs from both the periplasm and the cytoplasm to control precise gene expression patterns in diverse environmental conditions.

**Importance:** Despite high rates of vaccination, Pertussis, a severe, highly contagious respiratory disease, caused by the bacterium *Bordetella pertussis*, has reemerged as a significant health threat. In *Bordetella pertussis* and the closely related species, *Bordetella bronchiseptica*, activity of the BvgAS two-component regulatory system is critical for colonization of the human respiratory tract and other mammalian hosts, respectively. Here we show that the cytoplasmic PAS domain of BvgS can function as an independent signal perception domain that is capable of integrating environmental signals that influence overall BvgS activity. Our work is significant as it reveals a critical, yet previously unrecognized role, for the PAS domain in the BvgAS phosphorelay and provides a greater understanding of virulence regulation in *Bordetella*.

## Introduction

Pertussis, also known as whooping cough, is a severe, highly contagious, reemerging respiratory disease (1). It is caused by the bacterium *Bordetella pertussis*, which infects only humans and appears to be capable of surviving only transiently outside the human respiratory tract (2). By contrast, the closely related species *Bordetella bronchiseptica* infects nearly all mammals and can survive long periods of time in *ex vivo* environments, including those with scarce nutrients such as filtered pond water and buffered saline (3, 4). Despite these differences, *B. pertussis* and *B. bronchiseptica* produce a nearly identical set of virulence factors that are regulated at the level of transcription by functionally interchangeable BvgAS Two-Component Regulatory Systems (TCS) (5). By activating and repressing at least four classes of genes, BvgAS controls transition of the bacteria between three different phenotypic phases: the Bvg^+^ phase, which is necessary for virulence in mammals (6), the Bvg^−^ phase, which, in *B. bronchiseptica*, is required for survival under nutrient-limiting conditions (4) and for interactions with non-mammalian hosts (7), and the Bvg^i^ phase, which has been hypothesized to be important for bacterial transmission (8).

The BvgAS system is composed of a hybrid sensor kinase, BvgS, and a response regulator, BvgA (9). BvgS contains tandem, N-terminal, periplasmic Venus Flytrap (VFT) domains, followed by a transmembrane domain, a cytoplasmic Per/Arnt/Sim (PAS) domain, a histidine kinase (HK) domain, a receiver (REC) domain, and a C-terminal, histidine phosphotransfer (Hpt) domain (Fig. 1A). BvgA contains an N-terminal REC domain and a C-terminal helix-turn-helix (HTH) DNA binding domain. When *Bordetella* are grown on Bordet-Gengou (BG) blood agar or in Stainer-Scholte (SS) broth at 37°C, BvgS autophosphorylates, and, via a His-Asp-His-Asp phosphorelay, trans-phosphorylates BvgA. BvgA~P forms homodimers capable of binding DNA and activating or repressing gene transcription (10). Under these conditions, BvgA~P levels are high and the bacteria express the Bvg^+^ phase, which is characterized by transcription of all known virulence-activated genes (*vag*s), including those encoding critical virulence factors (11). When *Bordetella* are grown at 25°C or at 37°C in the presence of millimolar concentrations of nicotinic acid (NA) or magnesium sulfate (MgSO_4_), the BvgS phosphorelay is reversed and BvgS dephosphorylates BvgA, resulting in the Bvg^−^ phase. The Bvg^−^ phase is characterized by the expression of genes that are repressed by BvgA~P (termed virulence-repressed genes (*vrg*s)) and lack of expression of *vag*s (11). In *B. bronchiseptica*, flagellar regulon genes, including the flagellin-encoding gene *flaA*, are *vrgs* (12). When *Bordetella* transition between the Bvg^−^ phase to the Bvg^+^ phase or when the bacteria are cultured in the presence of low concentrations of NA or MgSO_4_, BvgA~P levels in the cell are low, resulting in the Bvg^+^ phase, which is characterized by repression of the *vrg*s, maximal expression of the intermediate phase gene *bipA*, and expression of *vag*s with promoters containing high affinity BvgA~P binding sites (such as *bvgAS* itself and *fhaB*, encoding the adhesin filamentous hemagglutinin (FHA)) but not *vag*s with promoters containing low affinity BvgA~P binding sites (such as the *ptx* operon, which encodes pertussis toxin and its export system) (10, 13, 14). The transition from Bvg^+^ phase to Bvg^−^ phase is termed phenotypic modulation.

**Figure 1.**
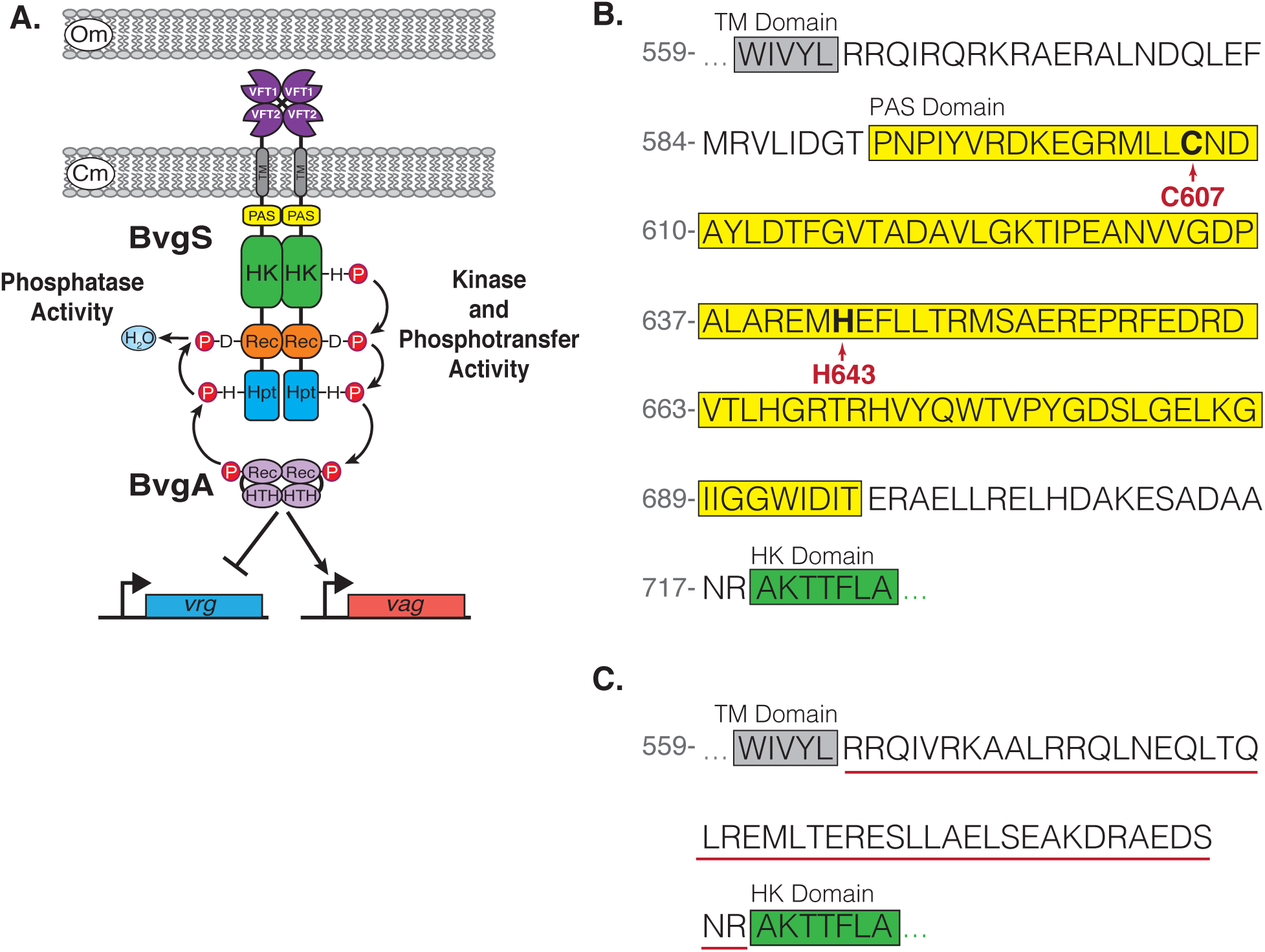
The sensor kinase, BvgS, has a unique PAS domain, which may play a role in signal perception. **(A)** Schematic of the two-component phosphor-relay system BvgAS. BvgS is an unorthodox His-Asp-His sensor kinase, comprised of tandem, periplasmic, venus fly-trap (VFT) domains followed by a transmembrane (TM) domain, a cytoplasmic Per/Arnt/Sim (PAS) domain, a histidine kinase (HK) domain, a receiver (REC) domain, and a histidine phosphotransfer (Hpt) domain. BvgA is a classical DNA-binding response regulator, comprised of a receiver (REC) domain and a helix-turn-helix (HTH) DNA-binding domain. When active, BvgS acts as a kinase and phosphorylates BvgA. The phosphorylated form of BvgA (BvgA~P) can bind DNA and activate or repress gene expression. When inactive, BvgS dephosphorylates and inactivates BvgA. **(B)** Amino acid sequence of the BvgS PAS domain (highlighted in yellow) and flanking domains (TM domain highlighted in grey and the HK domain highlighted in green). Red arrows highlight the highly conserved cysteine (C607) and histidine (His643) residues of the BvgS PAS domain that were targeted for mutagenesis. Grey numbers correspond to amino acid position in BvgS. **(C)** Schematic of the amino acid sequence of the alpha-helical linker (underlined in red) that was used to replace the BvgS PAS domain in the strain BvgS*_Ec_*. The amino acid sequence of the N-terminal TM domain is highlighted in grey and the C-terminal HK domain is highlighted in green. Grey number corresponds to amino acid position in BvgS.

Although the BvgAS phosphorelay and transcriptional control of the BvgAS regulon are well characterized, the natural signals that impact BvgS activity and the mechanism by which these signals are perceived and integrated by this system are unknown. Of BvgS’s three putative sensory perception domains, the periplasmic VFTs have been studied most. Structure-function analyses support a model in which VFT2 (the domain closest to the membrane) is in a closed conformation in the absence of modulating signals, and that this conformation renders BvgS active (15, 16). Hence the ‘default’ state for BvgS is one in which it is an active kinase. Binding of NA to VFT2 alters its conformation and this change is apparently transduced across the membrane, causing BvgS to reverse the phosphorelay and dephosphorylate BvgA, resulting in a shift to the Bvg^−^ phase (17).

The role of the PAS domain in regulating BvgS activity is less understood. PAS domains are abundant in prokaryotic signal transduction proteins and have a conserved structure, which includes a core binding pocket composed of a central, five-stranded antiparallel beta sheet and flanking alpha helices (18). Despite their conserved structure, PAS domains can perceive a multitude of stimuli, either by directly binding a signal molecule or by sensing with the assistance of a co-factor, in addition to promoting protein-protein interactions that mediate signal transduction (19–22). The PAS domain of BvgS contains two perception motifs that are critical for PAS domain function in other regulatory proteins; a quinone binding motif [aliphatic-(X)_3_-H-(X)_2,3_-(L/T/S)] and a highly conserved cysteine residue (C607) (23, 24). *In vitro* studies with the cytoplasmic portion of BvgS from *B. pertussis* showed that kinase activity was decreased in the presence of an oxidized ubiquinone analog and that the histidine in the quinone binding motif (H643) contributed to this effect (23). Jacob-Dubuisson and colleagues hypothesized that H643 may be involved in binding heme, but were unable to detect heme, or other cofactors, bound to the BvgS PAS domain (25). Their characterization of an H643A mutant of *B. pertussis* showed no defect in activation of the *ptx* promoter under aerobic conditions, but, instead, a defect in responding to NA and MgSO_4_. This group also investigated the importance of the conserved cysteine (C607) and found that a C607A mutant was similarly defective only in its ability to modulate in response to NA (25). Approximately one-third of BvgS homologs contain coiled-coil domains instead of PAS domains. Jacob-Dubuisson and colleagues showed that a *B. pertussis* mutant in which the BvgS PAS domain was replaced with the coiled-coil domain from the BvgS homolog of *Enterobacter cloacae* was similar to wild-type bacteria in its ability to activate *ptx* expression and to respond to NA (26). Taking these and other observations together, this group concluded that the role of the BvgS PAS domain is primarily to transduce signals from the VFTs to the kinase domain and that it may not function as a sensory perception domain to sense signals on its own.

Our laboratory has focused primarily on studying *B. bronchiseptica* because its broad host range allows us to study the role of *Bordetella* virulence factors and their regulation using natural host animal models (27, 28). We were motivated to investigate the role of the BvgS PAS domain during respiratory infection by our recent observation that maintenance of BvgS kinase activity in the lower respiratory tract (LRT) requires another TCS called PlrSR (29). PlrSR is a member of the NtrYX family of TCS, members of which are involved in sensing and responding to changes in redox status and anaerobiosis (30, 31). We hypothesized that, in the LRT, PlrSR controls expression of genes that contribute to electron transport-coupled oxidative phosphorylation and that deletion of *plrS* results in an altered redox state at the cytoplasmic membrane that is sensed by BvgS, perhaps via the PAS domain, resulting in its inactivation (i.e., converting it from a kinase to a phosphatase). In this study, we constructed *B. bronchiseptica* mutants with amino acid substitutions in the BvgS PAS domain and characterized them *in vitro* and *in vivo* to test this hypothesis.

## Results

### Mutants used in this study

To investigate the role of the BvgS PAS domain during infection, and to facilitate comparison of our results with those obtained with *B. pertussis*, we constructed mutant derivatives of *B. bronchiseptica* strain RB50 containing the same mutations as those that were present in the *B. pertussis* BPSM strains studied by Jacob-Dubuisson and colleagues (25, 26). Specifically, we constructed strains producing BvgS proteins in which H643 and C607 were replaced by alanine, and a strain in which the amino acids corresponding to the PAS domain were replaced with the analogous 47 amino acid (aa) region from the BvgS homolog of *Enterobacter cloacae* (Fig. 1B&C). We also constructed a strain in which BvgS C607 was replaced with the isosteric aa serine. As controls, we included the Bvg^+^ and Bvg^−^ phase-locked strains RB53 (aka BvgS^c^) and RB54 (aka Δ*bvgS*), respectively.

### Contribution of the PAS domain in regulating BvgS activity *in vitro*

To assess BvgS activity and sensitivity to modulating chemicals in our PAS domain mutants, we used two different *gfp* reporters; one containing the *fhaB* promoter (P*_fhaB_*), which contains high-affinity BvgA~P binding sites and therefore requires only very low levels of BvgA~P for activation, and one containing the *B. pertussis ptxA* promoter (P*_ptxA_*), which contains low-affinity BvgA~P binding sites and requires high levels of BvgA~P for activation (10, 13, 32, 33). The lower-affinity BvgA~P binding sites allow the P*_ptxA_-gfp* fusion to discern a broader range of BvgA~P levels (and hence BvgS kinase activities) than the P*_fhaB_*-*gfp* fusion. As expected, both promoters were activated in wild-type bacteria (WT), but not in the Δ*bvgS* mutant, when grown in SS broth at 37°C (Bvg^+^ phase conditions). P*_fhaB_-gfp* expression was similar in the strain containing the *bvgS-*C3 mutation (BvgS^c^) to that in WT, while P*_ptxA_-gfp* expression was much higher in BvgS^c^ than in WT (Fig. 2A&B), indicating that BvgS kinase activity is greater in BvgS^c^ than in WT when grown in SS at 37°C. When the bacteria were grown in medium containing 4 mM NA or 50 mM MgSO_4_ (Bvg^−^ phase conditions), P*_fhaB_-gfp* and P*_ptxA_-gfp* expression was very low in the wild-type and Δ*bvgS* strains and remained high in BvgS^c^, demonstrating the requirement of BvgS for activation of these promoters and the insensitivity of BvgS to NA and MgSO_4_ in BvgS^C^ (Fig. 2A&B).

**Figure 2.**
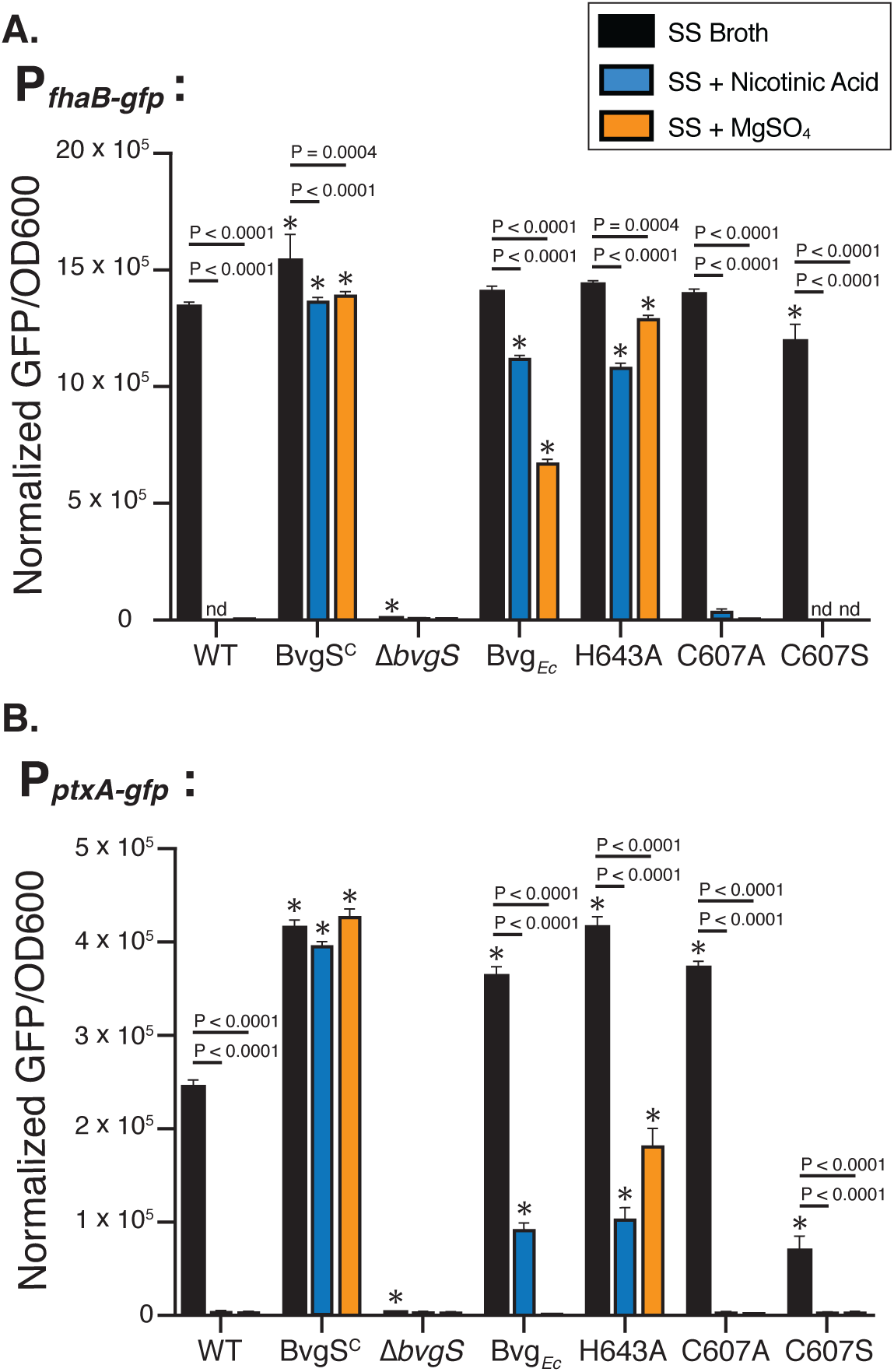
In *B. bronchiseptica*, the PAS domain influences BvgS kinase activity and sensitivity to chemical modulation *in vitro*. Transcriptional analysis of P*_ptxA_* and P*_fhaB_*activity in wild-type (WT), BvgS^C^, Δ*bvgS*, BvgS*_Ec_*, H643A, C607A, and C607S strains of *B. bronchiseptica* using the *gfp* based reporters. All strains were cultured in Stainer Scholte (SS) broth (Bvg^+^ growth conditions, black bars), in SS with 4mM nicotinic acid (Bvg^−^ growth conditions, blue bars), or in SS with 50mM MgSO_4_ (Bvg^−^ growth conditions, orange bars) at 37 °C with agitation. **(A)** Activity of the high-affinity BvgA~P P*_fhaB_* promoter was determined using the P*_fhaB_-gfp* reporter and **(B)** activity of the low-affinity BvgA~P P*_ptxA_* promoter was determined using the P*_ptxA_-gfp* reporter. Colored bars represent the mean normalized GFP fluorescence per cell (GFP/OD_600_) calculated from triplicate experiments. Error bars represent the standard deviation of normalized GFP/OD_600_ for each data set. Statistical significance (Two-Way ANOVA and Tukey’s multiple comparisons test, P < 0.05) of values for each strain compared to WT under each growth condition is indicated with * (P < 0.05) and statistically significance comparisons of values for a single strain cultured in each growth condition is indicated with a specific P value.

The “PASless” mutant (BvgS*_Ec_*, the strain in which the BvgS PAS domain was replaced with the analogous sequence from the BvgS homolog in *E. cloacae*) activated P*_fhaB_*-*gfp* to the same level as WT and P*_ptxA_-gfp* to a greater level than WT when grown under Bvg^+^ phase conditions (Fig. 2A), suggesting that the PASless BvgS is a more efficient kinase than wild-type BvgS (Fig. 2B). Expression of P*_ptxA_-gfp* and P*_fhaB_*-*gfp* was significantly decreased when BvgS*_Ec_* was cultured in the presence of NA or MgSO_4_ compared to growth in the absence of these chemical modulators (Fig. 2A&B), indicating that the PASless BvgS can sense MgSO_4_ and NA.

The H643A strain activated P*_fhaB_* and P*_ptxA_* similarly to BvgS^C^ and BvgS*_Ec_* when cultured in SS medium at 37°C, indicating that the H643A substitution does not disrupt BvgS’s ability to function as a kinase, and indeed makes it a more efficient kinase, *in vitro* (Fig. 2A&B). When grown in medium containing NA or MgSO_4_, P*_fhaB_*-*gfp* and P*_ptxA_-gfp* expression was significantly reduced in the H643A mutant compared to growth in the absence of NA and MgSO_4_, indicating that the H643A substitution does not affect BvgS’s ability to detect these chemicals.

Substitution of C607 with alanine or serine affected BvgS activity differently, despite both substitutions ablating the reactivity of the cysteine. The C607A mutation resulted in increased BvgS kinase activity (increased P*_ptxA_-gfp* expression) under Bvg^+^ phase conditions (Fig. 2A&B). By contrast, the C607S substitution resulted in decreased kinase activity (decreased P*_ptxA_-gfp* expression) under Bvg^+^ phase conditions. Neither mutation affected BvgS’s ability to respond to NA or MgSO_4_ compared to WT (Fig. 2A&B).

In all of the BvgS PAS domain mutants, BvgS remained responsive to MgSO_4_ and NA, indicating that the PAS domain is not required for sensing MgSO_4_ or NA. However, the level of BvgS activity in the presence of NA and MgSO_4_ differed dramatically between strains, suggesting that the PAS domain of BvgS does play a role in transducing signals sensed by the VFTs to the cytoplasmic HK, REC and HPT domains, which is consistent with previous reports (25).

### Characterization of pGFLIP-P*_flaA_* and pBam reporters in BvgS PAS domain mutants *in vitro*

To investigate the role of the PAS domain *in vivo*, we sought to measure bacterial burden and BvgS activity in the respiratory tract. To measure BvgS activity *in vivo*, we used two reporters that we have used previously (29, 34, 35). pGFLIP-P*_flaA_* is a recombinase-based reporter that indicates whether the bacteria have expressed the *flaA* gene, and hence the Bvg^−^ phase, at any time during the course of the experiment. Expression of *flaA* results in a conversion of the bacteria from GFP^+^ and Km^r^ to GFP^−^ and Km^s^ (34). By contrast, the pBam reporter provides an indication of whether the bacteria are in the Bvg^−^ phase at the time of plating on BG-blood agar. For wild-type *B. bronchiseptica* RB50, approximately 1 – 5% of bacteria containing the pBam plasmid will form large colonies (LCPs, for large colony phenotype) when plated and grown under Bvg^+^ phase conditions after growth under Bvg^−^ phase conditions (35). Although this reporter vastly under-estimates the number of bacteria that are in the Bvg^−^ phase at the time of sampling, it has proven useful in discerning differences in BvgAS activity during growth *in vitro* and in the lower respiratory tract (LRT) (29, 35).

We first characterized the use of pGFLIP-P*_flaA_*and pBam in our PAS domain mutants *in vitro*. When grown overnight in SS broth at 37°C (Bvg^+^ phase conditions), all strains maintained GFP fluorescence and formed few LCPs, indicating the bacteria did not modulate under these conditions (Fig. 3A-D). Consistent with our previous results (29), when grown with either 4 mM NA or 50 mM MgSO_4_ (Bvg^−^ phase conditions), nearly 100% of WT lost GFP fluorescence and ~2-10% formed LCPs, indicating that a majority of the cells had modulated in response to these chemicals and were in the Bvg^−^ phase at the time of plating (Fig. 3 A-D). For the BvgS*_Ec_*, C607A, and C607S strains, a majority of bacteria recovered after growth with NA and MgSO_4_ had also lost GFP fluorescence, indicating that they had modulated to the Bvg^−^ phase like WT. By contrast, very few GFP^−^ colonies were recovered for the H643A mutant, demonstrating the inability of this strain to modulate to the Bvg^−^ phase in response to NA and MgSO_4_. Taken with the P*_fhaB_*-*gfp* and P*_ptxA_*-*gfp* fusion data, these results indicate that although BvgS with the H643A substitution is able to sense NA and MgSO_4_, it does not dephosphorylate BvgA~P as completely as wild-type BvgS such that P*_flaA_* is expressed.

**Figure 3.**
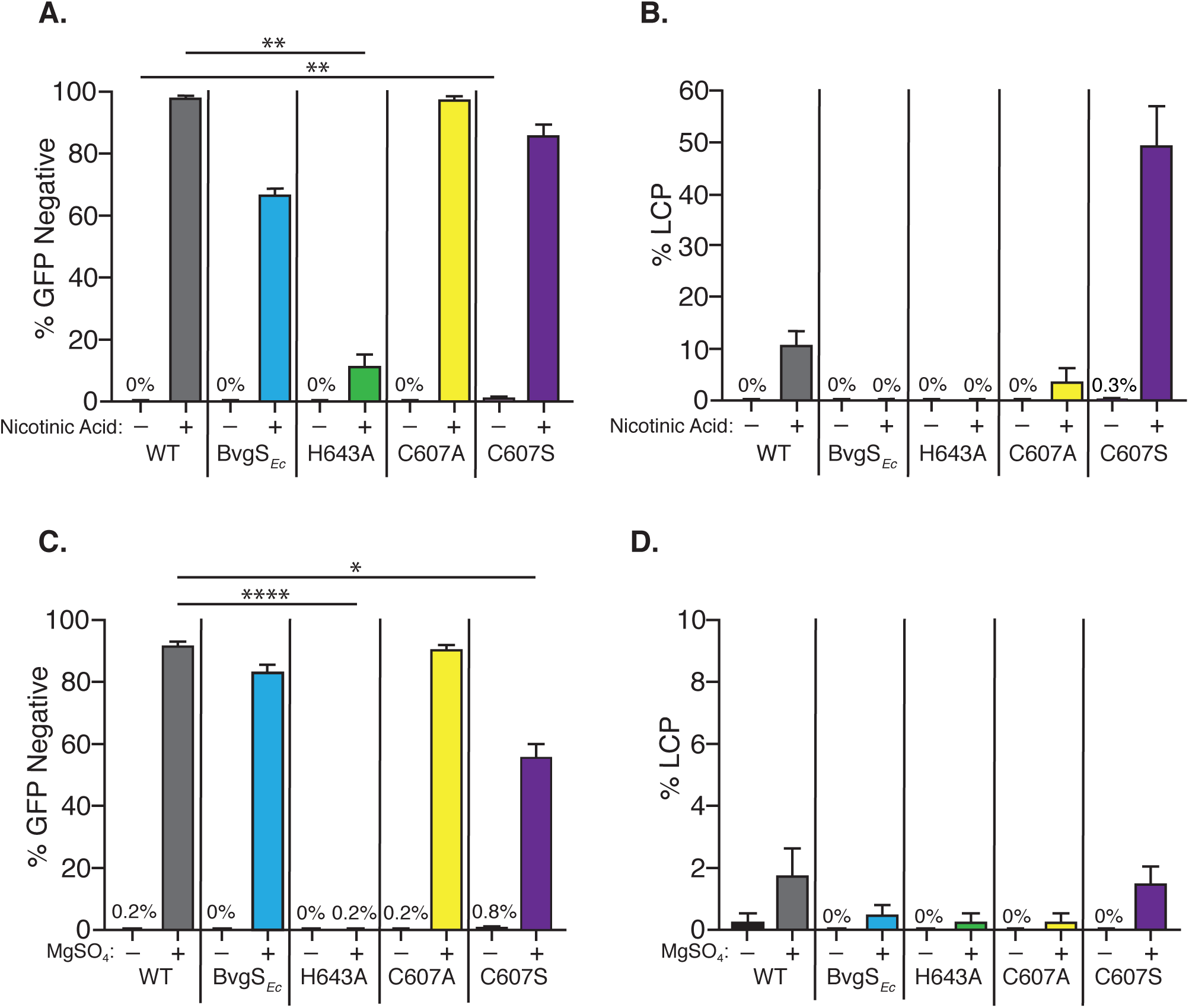
The PAS domain is important for BvgS inactivation in response to known modulatory chemicals *in vitro*. *In vitro* analysis of wild-type (WT), BvgS*_Ec_*, H643A, C607A, and C607S modulation in response to the chemical signals nicotinic acid (NA) and magnesium sulfate (MgSO_4_). All strains contain the plasmid reporters pGFLIP-P*_flaA_* and pBam. Strains were cultured under Bvg^+^ (SS broth) and under two different Bvg^−^ phase growth conditions (SS broth with 4 mM NA or 50 mM MgSO_4_) at 37 °C for ~18 h with agitation and subsequently plated on BG-blood solid agar to enumerate the percentage of GFP^−^ colonies **(A and C)** and LCPs **(B and D)**. NA was used as the modulatory stimulus in **(A)** and **(B)**. MgSO_4_ was used as the modulatory stimulus in **(C)** and **(D)**. GFP negativity indicates bacterial modulation and loss of BvgS kinase activity. LCPs indicate bacteria present in the Bvg^−^ phase when plated. Statistical significance (Kruskal-Wallis test and Dunn’s multiple comparisons test, P < 0.05) is indicated with * (P < 0.05), ** (P < 0.01), and **** (P < 0.0001).

In addition to indicating whether the bacteria are phenotypically Bvg^−^ phase at the time of plating, the pBam reporter provides an indication of how quickly BvgS transitions from phosphatase activity to kinase activity, because LCPs are formed by delayed or defective positive autoregulation of the *bvgAS* promoter (35). Although the differences were not statistically significant, the proportion of LCPs for the BvgS*_Ec_*, H643A, and C607A mutants after growth in MgSO_4_, and for the BvgS*_Ec_* and H643A mutants after growth in NA, were lower than for WT, consistent with BvgS in these strains having increased kinase activity (and hence increased positive autoregulation) compared to wild-type BvgS, as suggested by the P*_ptxA_*-*gfp* data (Fig. 2B). By contrast, the C607S mutant formed dramatically more LCPs than WT bacteria after growth in NA, consistent with this BvgS protein having decreased kinase activity compared to wild-type BvgS, which is also consistent with the P*_ptxA_*-*gfp* data. Altogether, these results indicate that the pGFLIP-P*_flaA_* and pBam reporters should be reliable indicators of BvgS activity in our PAS domain mutants *in vivo*.

### The PAS domain is not required for BvgS kinase activity in the LRT in PlrS^WT^ bacteria

One hypothesis for how PlrSR influences BvgS activity when the bacteria are in the LRT is that the BvgS PAS domain senses an activating signal that requires PlrSR and is needed for maintenance of BvgS kinase activity (Fig. S1A). This signal would be absent in a Δ*plrS* mutant leading to BvgS inactivation (Fig. S1B). If true, we predicted that PASless BvgS in BvgS*_Ec_* would be unable to sense the PlrSR-dependent activating signal and hence BvgS would fail to function as a kinase in bacteria with wild-type PlrSR (as well as in Δ*plrS* bacteria). To test this hypothesis, we inoculated mice intranasally with WT and BvgS*_Ec_* containing pGFLIP-P*_flaA_* and pBam and determined the number of colony forming units (CFU), GFP fluorescence, and LCP formation of bacteria recovered from the nasal cavity (NC) and right lung lobes at 0, 1, and 3 days post-inoculation. The BvgS*_Ec_* mutant colonized the NC and lungs similar to WT and showed no evidence of modulation (no conversion to GFP^−^ and few, if any, LCPs) (Fig. 4A-C), indicating that PASless BvgS had maintained kinase activity throughout the infection. Therefore, either our initial hypothesis is incorrect or the PASless BvgS protein no longer requires a PlrSR-dependent activating signal to initiate kinase activity in the LRT.

**Figure 4.**
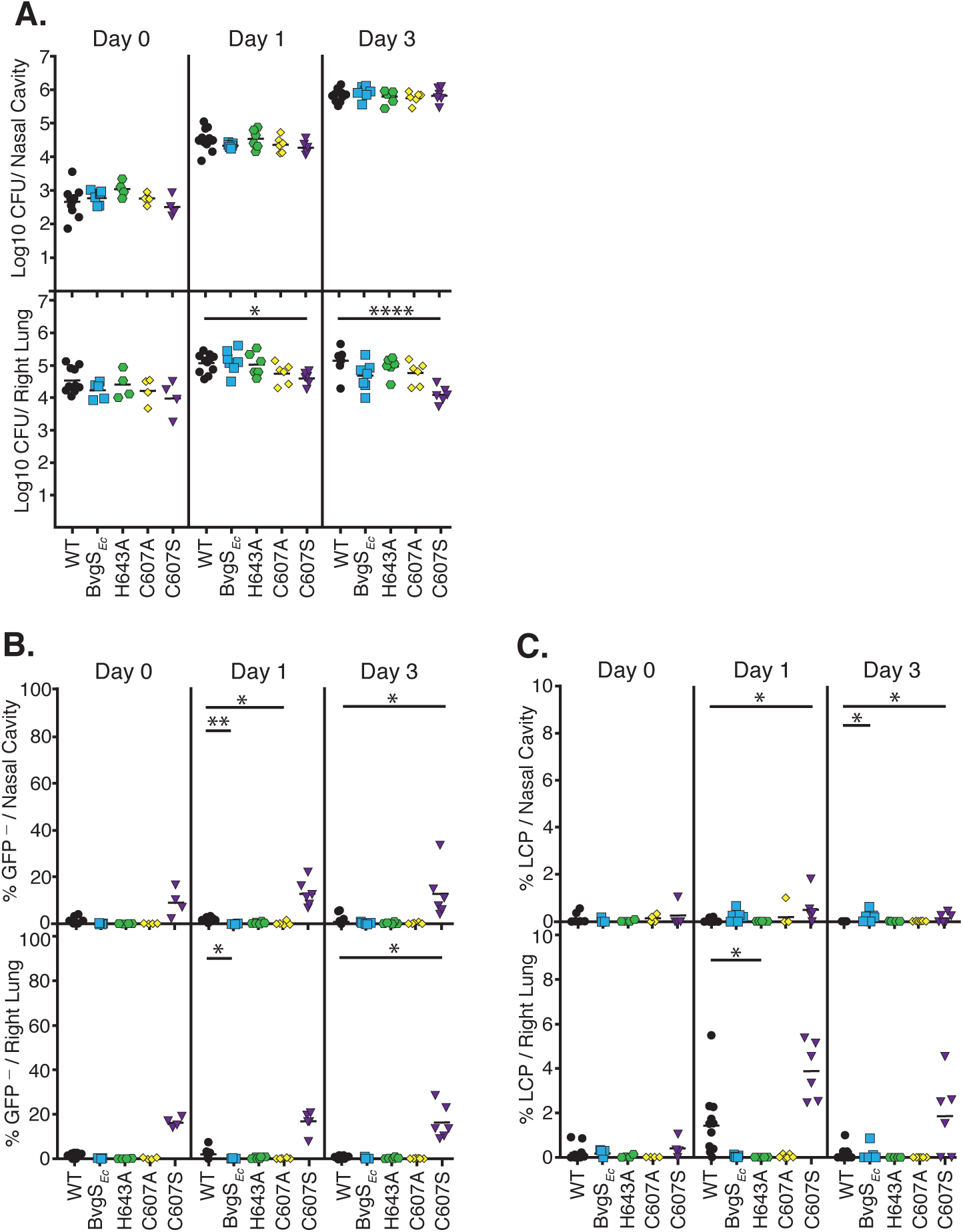
A conserved cysteine residue in the PAS domain is important for maintaining BvgS kinase activity during respiratory tract colonization of mice. **(A)** Colonization of the nasal cavity (top) and right lung (bottom) tissue in female BALB/cJ mice by wild type (WT; black circles), BvgS*_Ec_* (blue squares), H643A (green hexagons), C607A (yellow diamonds), and C607S (purple triangles) strains on days 0, 1, and 3 post-inoculation. All strains contain the plasmid reporters, pGFLIP-P*_flaA_*and pBam. Mice were inoculated with 7.5 ×10^4^ *cfu* via the external nares. Each symbol represents a single animal, with the mean colonization depicted as short horizontal bars. **(B)** Percentage of GFP negative (% GFP^−^) bacteria recovered from the nasal cavity (top) and right lung (bottom). GFP negativity indicates bacterial modulation and loss of BvgS kinase activity. **(C)** Percentage of large colony phenotype (LCP) producing bacteria recovered from the nasal cavity (top) and right lung (bottom). LCPs indicate bacteria present in the Bvg^−^ phase when recovered. Statistical significance (Kruskal-Wallis test and Dunn’s multiple comparisons test, P < 0.05) is indicated with * (P < 0.05), ** (P < 0.01), and **** (P < 0.0001).

### The C607S substitution hinders BvgS kinase activity and bacterial colonization in the lower respiratory tract (LRT) of mice

We also inoculated mice intranasally with WT, H643A, C607A, and C607S containing pGFLIP-P*_flaA_* and pBam to determine if the conserved histidine or cysteine residues in the PAS domain contribute to maintenance of BvgS kinase activity *in vivo*. While the H643A and C607A mutants colonized the NC and lungs similar to WT and showed no evidence of modulation (no conversion to GFP^−^ and few, if any, LCPs) (Fig. 4A-C), the C607S mutant was significantly defective for colonization in the lungs and a small proportion of the C607S bacteria recovered from the NC and lungs had modulated, as indicated by the loss of GFP and development of LCPs (Fig. 4B&C). These data are consistent with the *in vitro* data suggesting that the H643A and C607A mutants have increased BvgS kinase activity while the C607S mutant has decreased BvgS kinase activity when grown under Bvg^+^ phase conditions. If these proteins function *in vivo* as they do *in vitro*, our data suggest that decreased BvgS kinase activity, but not increased BvgS kinase activity, is detrimental to the ability of the bacteria to colonize the LRT, which is consistent with previous studies that demonstrate the necessity of the Bvg^+^ phase for mammalian respiratory tract infection (6, 36).

### A role for the BvgS PAS domain is revealed in the absence of PlrS in the LRT

A second hypothesis for how PlrSR influences BvgS activity when the bacteria are in the LRT is that PlrSR prevents the generation of a signal that would inhibit BvgS kinase activity (Fig. S1C). In a Δ*plrS* mutant, this inactivating signal would accumulate resulting in loss of BvgS kinase activity (Fig. S1D). If true, and if that signal is sensed by the BvgS PAS domain, then PASless BvgS, and potentially some of the PAS domain point mutants, would be insensitive to the inactivating signal and remain active in a Δ*plrS* mutant. To test this hypothesis, we deleted *plrS* in the BvgS PAS domain mutants and used the pGFLIP-P*_flaA_* and pBam reporters to evaluate BvgS activity in these strains *in vivo*. Similar to and as previously shown for the Δ*plrS* mutant (29), all strains containing the PAS domain mutations in addition to the Δ*plrS* mutation were severely defective for survival in the LRT, but colonized the NC similarly to WT (Fig. 5A). Similar to the parental Δ*plrS* strain, the C607A Δ*plrS* and C607S Δ*plrS* double mutants showed evidence of modulation in the LRT, with a large percentage of GFP^−^ bacteria recovered from the lungs at days 1 and 3 post-inoculation (Fig. 5B). A small proportion of bacteria of these same strains also modulated in the NC. These data suggest that C607 is not involved in the modulation of BvgS that occurs in the LRT in the absence of PlrS. Interestingly, despite the large number of GFP^−^ CFU recovered for the C607A Δ*plrS* mutant, very few LCPs were generated (Fig. 5C), suggesting that, in this strain, BvgS functions more efficiently as a kinase than wild-type BvgS once modulatory pressure is removed, consistent with the *in vitro* data.

**Figure 5.**
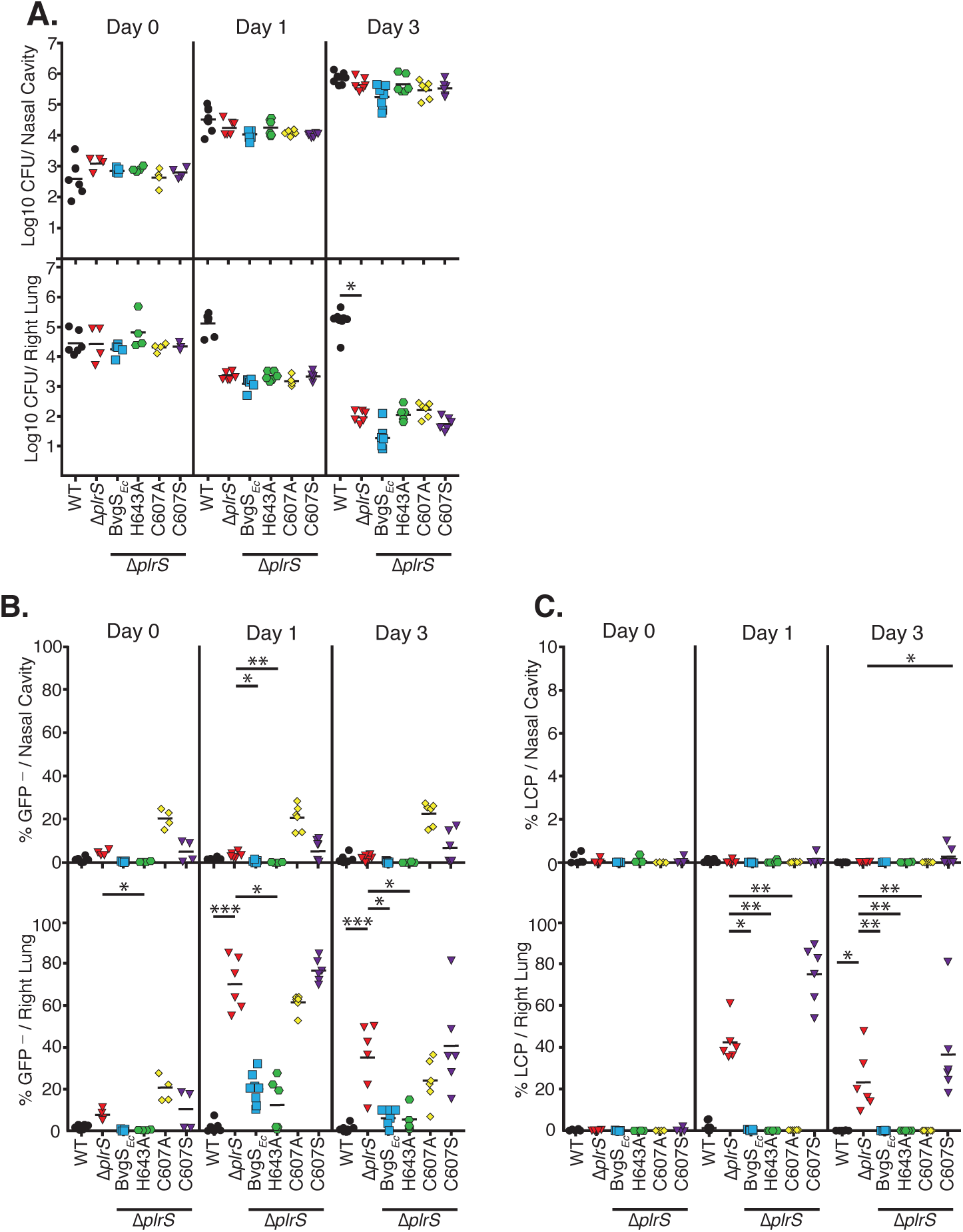
The PAS domain is required for BvgS modulation in response to PlrSR dysfunction during LRT colonization. **(A)** Colonization of the nasal cavity (top) and right lung (bottom) tissue in female BALB/cJ mice by wild type (WT; black circles), Δ*plrS* (red triangles), Δ*plrS*-BvgS*_Ec_* (blue squares), Δ*plrS*-H643A (green hexagons), Δ*plrS*-C607A (yellow diamonds), and Δ*plrS*-C607S (purple triangles) strains on days 0, 1, and 3 post-inoculation. All strains contain the plasmid reporters, pGFLIP-P*_flaA_*and pBam. Mice were inoculated with 7.5 ×10^4^ *cfu* via the external nares. Each symbol represents a single animal, with the mean colonization depicted as short horizontal bars. Homogenate from each organ was used to determine bacterial burden shown in graph A and *in vivo* bacterial modulation shown in graphs B and C. **(B)** Percentage of GFP negative (% GFP^−^) bacteria recovered from the nasal cavity (top) and right lung (bottom). GFP negativity indicates bacterial modulation and loss of BvgS kinase activity. **(C)** Percentage of large colony phenotype (LCP) producing bacteria recovered from the nasal cavity (top) and right lung (bottom). LCPs indicate bacteria present in the Bvg^−^ phase when recovered. Statistical significance (Kruskal-Wallis test and Dunn’s multiple comparisons test, P < 0.05) is indicated with * (P < 0.05), ** (P < 0.01), and *** (P < 0.001).

For the BvgS*_Ec_* Δ*plrS* and H643A Δ*plrS* mutants, and in contrast to the Δ*plrS* mutant, very few GFP^−^ colonies and few, if any, LCPs were recovered from the LRT (Fig. 5B&C), indicating that BvgS remained active in these strains, supporting the hypothesis that the PAS domain senses a negative signal that accumulates in the absence of PlrS in the LRT. *In vitro*, both of these mutant BvgS proteins were able to sense MgSO_4_ and NA, however, only the BvgS*_Ec_* mutant was capable of modulating to the Bvg^−^ phase sufficiently to activate P*_flaA_* in response to both chemicals. The fact that the BvgS*_Ec_* Δ*plrS* mutant did not activate P*_flaA_ in vivo* indicates that the modulating signal sensed by BvgS in the *ΔplrS* mutant background in the LRT is different, or is sensed differently, than NA or MgSO_4_ *in vitro*. These results suggest that the PAS domain, rather than the VFTs, is responsible for sensing the modulating signal that occurs in the LRT in the absence of PlrS. These results are the first to implicate the PAS domain of BvgS as an independent perception domain capable of influencing the activity of BvgS and support the hypothesis that the BvgS PAS domain may sense an inactivating signal that is regulated by PlrSR.

## Discussion

The fact that BvgS contains three putative signal perception domains and therefore has the potential to integrate multiple distinct environmental signals has been known for 20 years (21, 37, 38). To date, however, roles only for the VFT domains have been identified (15–17). Studies investigating the purpose of the PAS domain have suggested a role only in transducing signals sensed by the VFTs to the cytoplasmic signaling domains (25, 26). Our data support this role for the PAS domain. However, by investigating BvgS signaling during mammalian respiratory tract colonization, we have discovered that the PAS domain can function as a signal perception domain that is capable of influencing BvgS activity independently from the VFTs.

Our results cannot definitively distinguish whether the PAS domain senses an activating signal that is dependent on PlrSR (Fig. S1A) or senses an inhibitory signal that is produced in the absence of PlrSR (Fig. S1C). Regardless, our data support an important role for the PAS domain in sensing a signal(s) independent of the VFT domains. Our *in vitro* transcriptional analyses, showed that both wild-type BvgS and PASless BvgS can sense NA and MgSO_4_, presumably via the VFT domains, and in response they sufficiently modulate the bacteria to the Bvg^−^ phase. *In vivo*, wild-type BvgS was also sensitive to a PlrS-dependent signal in the LRT (it modulated in the Δ*plrS* strain), however, PASless BvgS was insensitive to this signal, indicating that this signal 1) is sensed differently than NA or MgSO_4_ (not via the VFTs) and 2) specifically requires the PAS domain for perception.

Similar to the PASless BvgS protein, BvgS with the H643A substitution was also insensitive to the PlrS-dependent signal *in vivo*; the H643A strain did not activate the P*_flaA_* promoter in the absence of PlrS in the LRT. Although this BvgS protein was capable of sensing NA and MgSO_4_ *in vitro*, it did not modulate sufficiently to de-repress P*_flaA_* under these conditions, and therefore, we cannot rule out that this strain did not modulate to some degree *in vivo* while in the LRT.

PlrSR is a member of the NtrYX family of TCS (39). Although some NtrYX family members are involved in nitrogen homeostasis, others, such as those in *Brucella abortus* and *Neisseria gonorrhoeae*, control genes involved in bacterial respiration (such as those encoding high affinity cytochrome oxidases) in response to low oxygen (30, 40). We hypothesize that PlrSR may similarly control the expression of genes that contribute to electron transport-coupled oxidative phosphorylation. Absence of appropriate cytochromes oxidases when the bacteria are in the LRT would likely alter the redox status of the electron transport chain components, such as ubiquinone, and would also likely prevent bacterial growth. Bock and Gross showed that BvgS kinase activity is inhibited by oxidized ubiquinone *in vitro* and that the H643 residue, which resides within a putative quinone binding motif in the BvgS PAS domain, contributes to BvgS’s sensitivity to this redox-active molecule (23). We are currently investigating whether PlrSR controls the expression of the multiple operons encoding high and low affinity cytochrome oxidases and, if so, their role in controlling BvgS activity in the LRT.

Although our data suggest that the conserved cysteine residue (C607) does not appear to contribute to the sensing capacity of the BvgS PAS domain (at least not while the bacteria are in the LRT), they do indicate that this residue is important for maintaining BvgS kinase activity. *In vitro*, substitution of alanine for C607 enhanced BvgS kinase activity, while the C607S substitution significantly decreased BvgS kinase activity. It was surprising that the C607A and C607S mutations had different effects on BvgS kinase activity. Although both substitutions abrogate the reactivity of the conserved cysteine, alanine and serine differ significantly by size and reactivity of their R-group. Alanine is a relatively small amino acid and nonreactive compared to cysteine and serine. Serine is capable of forming hydrogen bonds within a protein and contains a similarly sized R-group compared to cysteine. It is possible, based on our data, that when BvgS is in an active kinase confirmation C607 is modified in a way that reduces the size of its side chain and that maintenance of a bulky R-group within the PAS domain, as would occur with a serine amino acid, is disruptive and inhibits BvgS activation. This disruption would not occur with the C607A substitution and we find that this mutation only enhances BvgS kinase activity. Another possibility is that formation of aberrant hydrogen bonds by the non-native serine residue may destabilize the protein conformation that promotes BvgS kinase activity and would, therefore, disrupt protein function. To fully understand the impact these amino acid substitutions have on BvgS, additional biochemical characterization of these modified BvgS proteins is required.

Overall, our results are generally consistent with those of Jacob-Dubuisson and colleagues that demonstrated a role for the BvgS PAS domain in regulating efficient BvgS signal transduction (25). These results are not surprising as BvgS from *B. bronchiseptica* and *B. pertussis* are functionally interchangeable and the PAS domain is highly conserved between species (only differing by 2 amino acid residues) (5, 41). We did, however, identify some nuanced differences in BvgS activity between *B. bronchiseptica* and *B. pertussis*. When assessing BvgS kinase activity, our results indicated that BvgS in *B. bronchiseptica* fails to fully activate when the bacteria are grown under laboratory conditions. We showed that replacement of some amino acids in the PAS domain enhanced BvgS kinase activity compared to WT, indicating that the PAS domain influences BvgS kinase activity in *B. bronchiseptica*. This enhancement of kinase activity was not observed in *B. pertussis* with similar substitutions (25, 26), suggesting that BvgS may already be maximally active *in vitro* and/or the PAS domain does not influence BvgS activation in *B. pertussis*. These data could explain why some strains of *B. pertussis* appear less sensitive to modulating signals *in vitro* (NA and MgSO_4_) compared to most *B. bronchiseptica* isolates (5). If BvgS from *B. pertussis* has higher kinase activity *in vitro*, it is likely that more modulatory signal would be required to inhibit BvgS kinase activity sufficiently to noticeably decrease *vag* expression. We also observed that modification of the BvgS PAS domain had different effects on BvgS’s responsiveness to NA and MgSO_4_ in each species. In *B. bronchiseptica*, BvgS with the H643A substitution remained responsive to NA and MgSO_4_, while, in *B. pertussis*, this substitution rendered BvgS insensitive to both chemicals (25). Similarly, the C607A substitution made BvgS from *B. pertussis* insensitive to modulation by NA, however, BvgS responsiveness was unchanged by this substitution in *B. bronchiseptica* (25). As the VFTs are implicated in sensing NA and MgSO_4_, the differences we observed are likely due to the significant variability of amino acid sequence encoding the BvgS VFT domains in *B. bronchiseptica* and *B. pertussis* (5, 41). While our data corroborate a role for the PAS domain in facilitating the transduction of signals though the BvgS protein, our results also highlight that it is important to consider how the VFT domains may influence the function of the PAS domain and how the synergy of all perception domains contribution to the ultimate output of BvgS activity.

The VFT domains and the PAS domain serve as independent points of signal integration within BvgS. It is well known that the modulatory signals NA and MgSO_4_ require the VFT domains for perception *in vitro* and based on our working model, the PAS domain is perhaps sensitive to the redox state of ubiquinone. The natural signals sensed by BvgS, however, remain unknown. By having multiple separate signaling domains, BvgS has the potential to control bacterial gene expression appropriately in many environments. For example, in the mammalian LRT BvgS kinase activity is essential for bacterial survival, therefore, it is likely that both the VFT domains and the PAS domain are in an ‘On’ or active conformation, thus permitting BvgS activity. In a Δ*plrS* mutant in the LRT, however, BvgS is inactive and the PAS domain is likely in an ‘Off’ or inactive conformation based on our results. This environment, although artificial, potentially mimics a naturally occurring environment in which the PAS domain can integrate a signal that overrides an activating input sensed by the VFT domains, indicating the possibility that the VFT domains and the PAS domain could obtain various combinations of conformations, rendering BvgS on or off depending on which domain’s influence dominates within the specific environment encountered by the bacteria. Interestingly, not all BvgS homologs have PAS domains, suggesting that bacteria with simplified BvgS proteins perhaps do not encounter such a diverse range of environments and, therefore, do not require BvgS to integrate multiple signals (26). The fact that a PAS domain in BvgS has been maintained in *Bordetella* is indicative of the critical role this domain plays in the regulation of this important TCS.

## Experimental Procedures

### Bacterial Strains, Plasmids, and Growth Conditions

All bacterial strains and plasmids used in this study are listed in Table 1. *Escherichia coli* strains were grown using Luria-Bertani (LB) broth or agar at 37°C. *B. bronchiseptica* strains were grown on Bordet-Gengou (BG) agar supplemented with 7.5% defibrinated sheep blood (Hemostat, Cat#: DSB1) or in Stainer-Scholte (SS) broth supplemented with SS supplement (REF) at 37°C (42). As needed, media was supplemented with ampicillin (Amp; 100 μg/mL; Sigma, Cat#: A0166), streptomycin (Sm; 20 μg/mL; Gold Biotechnology, Cat#: S-150-50), gentamicin (Gm; 30 μg/mL; Gold Biotechnology, Cat#: G-400-25), kanamycin (Km; *E.coli* at 50 μg/mL or *B. bronchiseptica* at 125 μg/mL; Gold Biotechnology, Cat#: K-120-25), or diaminopimelic acid (DAP; 300 μg/mL; VWR, Cat#: AAB22391-06).

### Construction of *B. bronchiseptica* BvgS PAS Domain Mutant Strains

DNA manipulation and cloning were performed according to standard methods (43). All restriction enzymes and DNA ligase were obtained from New England BioLabs. High-fidelity PFU Ultra II DNA polymerase (Agilent, Cat#: 600670) was used for the construction of all allelic exchange vectors used in this study. GoTaq DNA polymerase (Promega, Cat#: M3001) was used for screening purposes during plasmid and strain construction. All enzymes were used according to the manufacturer’s instructions. The primers used in this study are listed in Table 2. All primers were synthesized by Eton Biosciences and purified using a standard desalting method.

To create the BvgS*_Ec_* mutation, primers MAB85-MAB90 were used to generate an approximately 1.2 kb DNA fragment, which contained a 141 bp sequence encoding an α-helical linker from the BvgS homolog in *Enterobacter cloacae* (GI: 915610361), flanked by homologous sequences (500 bp each) corresponding to the regions 5’ and 3’ to the PAS domain of the *B.b. bvgS* gene. Individual point mutations in *bvgS* (H643A, C607A, and C607S) were generated by constructing allelic exchange plasmids containing an approximately 1.0 kb DNA fragment of the *bvgS* sequence, centered on the PAS domain, with exact homology to *bvgS* except for the desired substitution and a unique restriction enzyme digestion site, used for screening. Primers used to generate the most 5’ and 3’ ends of the DNA fragments for each mutant were designed to include a BamHI restriction site at the 5’ end and a SacI restriction at the 3’ end, which were used to clone the DNA fragments into the allelic exchange plasmid pEG7S (4). Standard methods were then used to deliver this plasmid to RB50 and Δ*plrS* strains by conjugation using RHO3 cells. Double homologous recombination events that introduced the desired mutation into the chromosome were selected for using 15% sucrose and screened for by restoration of BvgS activity indicated by the presence of hemolysis on BG-blood agar or through restriction digestion using the unique restriction enzyme site introduced for point mutants. Sanger sequencing was used to verify each mutation in *bvgS*. The pGFLIP-P*_flaA_* and pBam reporters were introduced into each mutant strain, using tri-parental mating with conjugative RHO3 or by homologous recombination, respectively. Introduction of each reporter on to the chromosome was verified using PCR.

### Construction of P*_ptxA-gfp_* and P*_fhaB-gfp_*Transcriptional Reporters

To construct a GFP transcriptional reporter for P*_ptxA_*, the *ptxA* promoter (position −5 to −220 upstream of *ptxA* coding sequence) was amplified by PCR from *B. pertussis* Tohama I. To construct a GFP transcriptional reporter for P*_fhaB_*, the *fhaB* promoter (position −5 to −254 upstream of *fhaB* coding sequence) was amplified by PCR from *B. bronchiseptica* RB50. PCR products were then digested and cloned into the pUC*gfp*MAB plasmid using EcoRI and HindIII restriction sites. The resulting plasmids were subsequently sequenced to verify that each promoter was correctly fused 5’ to the S12 RBS and *gfp*. Each reporter plasmid was then introduced into WT, BvgS^C^, Δ*bvgS*, BvgS*_Ec_*, H643A, C607A, and C607S strains using tri-parental mating with conjugative RHO3. A strain of RB50 was also engineered to contain the pUC*gfp*MAB plasmid without a promoter driving *gfp* expression as a control (RB50::pUC*gfp*MAB-EV). Reporter delivery to the Tn7 site was confirmed by PCR in each strain.

### Transcription of P*_ptxA-gfp_* and P*_fhaB-gfp_*Reporters *In Vitro*

To analyze expression of the P*_ptxA-gfp_*and P*_fhaB-gfp_* transcriptional reporters under Bvg^+^ phase conditions, strains were grown on BG-blood agar supplemented with kanamycin at 37 °C until colonies formed. Approximately 3-4 colonies were used to inoculate liquid SS media containing streptomycin and kanamycin, with or without 4mM nicotinic acid (NA) or 50mM MgSO_4_, depending on the experimental conditions tested. All cultures were incubated overnight for ~18 h at 37 °C on a roller. Each overnight culture was then sub-cultured to an OD of 0.3 per milliliter into fresh liquid SS media containing streptomycin and kanamycin, with or without 4mM nicotinic acid (NA) or 50mM MgSO_4_, depending on the experimental conditions tested. Subcultures were incubated for ~4 h at 37°C while rotating. Subsequently, one-hundred microliters of each culture were transferred to a well of a 96-well, flat bottom, polystyrene plate (Greiner Bio-One CellStar, Cat#: 655180), and the OD600 and RFU at excitation A485 and emission A535 (to detect GFP) were measured on a Perkin-Elmer Victor3 1420 Multi-label plate reader. Total GFP expression was normalized by initial subtraction of background fluorescence, as determined from the RB50::pUC*gfp*MAB-EV strain, and normalization to the OD600 for each sample. Cultures were prepared and tested in duplicate and three experimental replicates were completed. Normalized GFP/OD600 values were averaged across all replicates.

### Characterization of pGFLIP-P*_flaA_* and pBam Reporters *In Vitro*

Strains containing pGFLIP-P*_flaA_* and pBam reporters were grown overnight in SS medium with streptomycin and gentamycin, with and without 50 mM MgSO4 or 4 mM nicotinic acid. After incubation at 37 °C for 24 h on a rotating incubator, an aliquot of cells from each experimental condition was diluted to a concentration of 3 × 10^3^ cells/mL with 1X PBS and plated on BG-blood agar in duplicate. Plates were incubated for 72 h at 37 °C. White light and GFP fluorescence images of each plate were captured using a G:Box Imaging System (Syngene). Total *cfu*, GFP positive colonies, and LCP colonies were enumerated using the colony analysis protocol of the GeneTools Software package (Syngene). The total number of GFP negative colonies was calculated by subtracting the number of GFP positive colonies from the total number of colonies isolated. The percent GFP negative colonies and percent LCP colonies were calculated based on the total number of colonies isolated. Each *in vitro* experimental condition was tested twice.

### Evaluation of *B. bronchiseptica* Modulation *In Vivo*

Bacteria were cultured overnight at 37 °C in SS media with streptomycin, kanamycin, and gentamicin to maintain bacteria in the Bvg^+^ phase (42). Six-week-old female BALB/c mice from Jackson Laboratories were inoculated intranasally with 7.5 × 10^4^ *cfu* of *B. bronchiseptica* strains (containing the pGFLIP-P*_flaA_* and pBam reporters) in 50 μL of 1 X DPBS. At each time point reported, right lung lobes and nasal cavity tissue were harvested and homogenized in 1 X DPBS using a mini-beadbeater. Dilutions of the tissue homogenates were then plated on BG-blood agar and grown at 37°C for 72 h. White light (to enumerate total *cfu* and LCPs) and blue light (to enumerate GFP^+^ colonies) images of each plate were captured using a G:Box Imaging System (Syngene). Total *cfu*, GFP positive colonies, and LCP colonies were enumerated using the colony analysis protocol of the GeneTools Software package (Syngene). The total number of GFP negative colonies was calculated by subtracting the number of GFP positive colonies from the total number of colonies isolated. The percent GFP negative colonies and percent LCP colonies were calculated by determining the ratio of GFP negative colonies or LCP colonies to the total number of colonies isolated.

## Ethics Statement

This study was carried out in strict accordance with the recommendations in the Guide for the Care and Use of Laboratory Animals of the National Institutes of Health (44). All animals were properly anesthetized for inoculations, monitored regularly, and euthanized when moribund. Efforts were made to minimize suffering. Our protocol was approved by the University of North Carolina Institutional Animal Care and Use Committee (Protocol ID: 16-246).

## Statistical Analysis

Statistical analysis was performed using Prism 8.0.1 software from GraphPad Software. Statistical significance was determined using a Two-Way ANOVA and Tukey’s multiple comparisons test (*gfp* transcriptional assays) or a Kruskal-Wallis test and Dunn’s multiple comparisons test (*in vitro* and *in vivo* modulation assay using pGFLIP-P*_flaA_* and pBam). P < 0.05 was considered significant. Figures were generated using Adobe Illustrator CS6 (Adobe Systems).

## Acknowledgments

We thank members of the Cotter lab for critical discussions and technical assistance. We thank Françoise Jacob-Dubuisson for providing the plasmid used in making the BvgS*_Ec_*strain. This work was supported by NIH grant R01 R01AI129541 to P.A.C.

## Author Contributions

M.A.B. and P.A.C. designed research; M.A.B. performed research; M.A.B. and P.A.C. analyzed data; and M.A.B. and P.A.C. wrote the paper.

The authors declare no conflict of interest.

---

**Supplemental Figure 1. The PAS domain of BvgS may sense an activating or inactivating signal in the LRT. (A)** Schematic depicting how the PAS domain may be required to sense a PlrS-dependent activating signal that is required to maintain BvgS kinase activity in the LRT. Genes encoding a protein(s) required for the generation of an activating signal are regulated by the PlrSR TCS. The BvgS PAS domain is required for sensing this activating this signal and BvgS can function as a kinase. **(B)** Without PlrS, generation of the activating signal does not occur and BvgS would be off. **(C)** Schematic depicting how the PAS domain may sense a signal that inactivates BvgS in the LRT. Normally, the activity of a protein(s) encoded by a gene(s) regulated by the PlrSR system would inhibit the generation of an inactivating signal. **(D)** However, without PlrS this inactivating signal can accumulate and inhibit BvgS kinase activity via the PAS domain of the protein.

